# Renal cancer cell-derived amphiregulin recruits mesenchymal stromal cells, induces their glycolytic switch, and promotes tumour growth

**DOI:** 10.1101/2025.10.14.681141

**Authors:** Piotr Popławski, Małgorzata Grzanka, Tomasz Skirecki, Grażyna Hoser, Kacper Bielak, Dominika Nowis, Helena Kossowska, Roksana Iwanicka-Nowicka, Marta Koblowska, Urszula Jankowska, Bożena Skupień-Rabian, Anna Burdzińska, Weronika Zarychta-Wiśniewska, Michał Pyzlak, Anna Litwiniuk, Magdalena Bandyszewska, Joanna Życka-Krzesińska, Joanna Bogusławska, Natalia Rusetska, Elżbieta Sarnowska, Beata Rybicka, Alex Białas, Katarzyna D. Arczewska, Michał Mączewski, Wojciech Bik, Leszek Pączek, Agnieszka Piekiełko-Witkowska

## Abstract

Mesenchymal stem/stromal cells (MSCs) are multipotent cells that support wound healing. Tumours, often called as wounds that do not heal, recruit MSCs, which in turn support tumour growth. The immunomodulatory MSCs properties are facilitated under hypoxic conditions, while tumours are often hypoxic and immune-suppressed. It is unclear how MSCs wound healing actions are prevented in tumours.

Here, we found that renal cancer cells secrete amphiregulin which induces MSCs recruitment and facilitates tumour growth in vivo. In MSCs, AREG triggers HIF1A degradation and transcriptional reprogramming, leading to glycolytic switch, increased migration and attenuation of immunoregulatory profile. In renal cancer cells, AREG stimulates expression of oncogenic proteins, including AKR1C3, leading to increased tumour growth and angiogenesis. To our knowledge, this is the first study showing that glycolytic switch in MSCs is induced by cancer.

Our study provides insight into how renal cancer cells attenuate MSCs wound pro-healing properties to facilitate tumour growth.

## Introduction

Mesenchymal stem/stromal cells (MSCs) are multipotent cells capable of differentiating into several cell type lineages. Their primary site of origin is the bone marrow from which they can be recruited to the various target sites and differentiate into other types of cells, including adipocytes, chondrocytes and osteocytes. Under physiological conditions, MSCs are recruited to injured sites to support wound healing. On the other hand, MSCs are also involved in severe pathological conditions, including fibrosis and cancer (Antoon et al., 2024; Qin et al., 2023).

Cancerous lesions are surrounded by rich tumour microenvironment (TME) formed by cellular and non-cellular components that facilitate survival of neoplastic cells and promote malignant progression. Tumours, which are often referred to as the ‘wounds that do not heal’, secrete chemokines that recruit MSCs, which in turn secrete cytokines and growth factors to promote angiogenesis, induce extracellular matrix (ECM) remodelling, apoptotic resistance and immune evasion (Antoon et al., 2024). MSCs can also differentiate into CAFs (cancer associated fibroblasts) that further support tumour growth (Sun and Yao, 2023). Compared to other TME components, MSCs represent relatively small percentage of TME cells; however, their impact is disproportionally large due to their direct effects on ECM remodelling and regulation of interactions between other cellular TME components (Sun and Yao, 2023). The pro-healing properties of MSCs are more enhanced under hypoxic conditions (Mathew et al., 2017; Riis et al., 2017; Y. Yang et al., 2022). At the same time, hypoxia is one of the key hallmarks of solid cancers (Z. Chen et al., 2023) which creates a puzzling conundrum of the mechanisms that prevent activation of the MSCs wound healing actions in tumours.

Renal cell cancer (RCC) is the most common kidney malignancy that affects 400,000 people worldwide annually. The vast majority of RCC diagnoses is classified as clear cell RCC (ccRCC), representing 75-80% of cases. Despite the progress in treatment of advanced ccRCC, including the introduction of immune-checkpoint inhibitors, metastatic RCC still remains incurable disease, leading to death of 180,000 patients per year (Makino et al., 2022). The key molecular abnormalities linked with ccRCC include inactivating mutations of VHL tumour suppressor that lead to the permanent activation of the hypoxia-induced pathways, induction of angiogenesis and metabolic reprogramming (Warburg effect) (Cancer Genome Atlas Research Network, 2013). One of the less understood ccRCC TME components are MSCs which are recruited to the RCC tumours and support cancerous progression. Specifically, co-culturing of MSCs with RCC cells augments migration and invasiveness of the latter (Hsiao et al., 2012) . MSCs can also release microvesicles that promote ccRCC growth and malignancy in vitro and in vivo (Du et al., 2014), while MSCs pre-stimulated by extracellular vesicles released by RCC stem cells enhance migration of renal cancer cells and formation of blood vessels (Lindoso et al., 2015). However, the exact mechanisms that govern MSCs recruitment to RCC tumours are poorly understood. In our recent study we identified AREG and fibronectin (FN1) as the components of ccRCC secretome that stimulated MSCs migration in vitro (Popławski et al., 2023). Here, we aimed to explore the impact of AREG and FN1 on the recruitment of MSCs in vivo. Surprisingly, we found that AREG acted bi-modally: as a secreted growth factor that induces MSCs migration, as well as autocrine factor that stimulates growth of ccRCC tumours. Moreover, our study provides insight into how ccRCC attenuates MSCs’ wound pro-healing properties to facilitate tumour growth.

## Material and Methods

ccRCC-derived cell lines: 786-O (cat. No. CRL-1932) and Caki-1 (cat. no. HTB-46) were purchased from ATCC and were cultured in accordance with manufacturer’s protocols.

### MSCs isolation

Bone marrow (BM) aspirates for MSCs isolation were obtained during routine orthopedic procedure, following approval from the Local Bioethics Committee (Approval No. KB/115/2016) and with written informed consent from patient. BM samples were mechanically disaggregated and passed through a 21G needle, then diluted 1:2 with PBS and layered over Histopaque-1077 (Sigma-Aldrich, St. Louis, MO, USA), followed by centrifugation at 400 × g for 30 min at RT. Mononuclear cells were collected from the interface, washed twice with PBS, and resuspended in low-glucose Dulbecco’s Modified Eagle Medium (Biowest, Riverside, MO, USA) supplemented with 10% fetal bovine serum (FBS), Glutamine-Penicillin–Streptomycin, and amphotericin B (all from Biowest, Riverside, MO, USA). BM-MSCs identity was verified by analysis of surface markers using BD Stemflow™ Human MSC Analysis Kit (BD Biosciences; CAT# 562245) and by confirmation of their differentiation ability into adipogenic, chondrogenic, and osteogenic lineages, as described previously (Zielniok et al., 2020). The cytometric and differentiation characteristic of MSCs is provided in **Supplementary Figure S1**.

Vectors pLenti-C-mGFP-P2A-Puro_AREG (Origene CAT#: RC203150L4), pLenti-C-mGFP-P2A-Puro_FN1 (Origene CAT#: RC212860L4), and empty vector pLenti-C-mGFP-P2A-Puro (CAT#: PS100093) were purchased from Origene (Rockville, MD).

### Cell line transduction

786-O cell line was transduced using lentiviral transduction as previously described (Winiarska et al., 2017). For lentivirus production, 1.5 × 10^6^ HEK-293T cells were seeded into 10 cm dishes 72 h prior to transfection in 10 ml of DMEM supplemented with 10% FBS (Invitrogen) and P/S solution (ThermoFisher, Waltham, MA, USA). The lentiviruses encoding the gene of interest (GOI) were introduced to the packaging HEK293T cells by calcium phosphate co-transfection with 8.6 μg of a GOI-containing vectors (pLenti-C-mGFP-P2A-Puro_AREG, pLenti-C-mGFP-P2A-Puro_FN1, or pLenti-C-mGFP-P2A-Puro) and components of 2^nd^ generation of packaging vectors: 8.6 μg of psPAX2 packaging vector and 5.5 μg of pMD2.G envelope vector (a kind gift of Professor Didier Trono; École Polytechnique Fédérale de Lausanne, Lausanne, Switzerland). Right before transfection HEK293T culture medium was exchanged to 5ml of fresh DMEM+ 10% FBS w/o P/S. The plasmids were resuspended in 450 μl of 300 mM CaCl_2_ and added dropwise to 450 μl of 2 × concentrated HEPES-buffered saline (280 mM NaCl, 20 mM HEPES, 1.5 mM Na2HPO_4_, 10 mM KCl and 12 mM D-glucose pH7.2) by vortexing. The precipitate was immediately added to the cell culture medium with gentle swirling. The medium was replaced with 6 ml of fresh DMEM+ 10% FBS+ P/S 16 h post transduction. Virus-containing supernatants were collected 48 h later, centrifuged for 5 min at 300 × g and filtered through 0.45 μm low-protein-binding filters (Merck Millipore, Darmstadt, Germany). The titer of viral particles was estimated using a Lenti-X p24 Rapid Titer Kit (Takara Bio) according to the manufacturer’s protocol. MOI was calculated by dividing the vector titer for the number of cells transduced. 786-O and Caki-1 cells were transduced using a multiplicity of infection (MOI) of 3. For initial selection, the transduced cells were cultured in 2.5 μg/ml puromycin (Gibco, ThermoFisher, Waltham, MA, USA) for 5 days, followed by sorting of fluorescent-positive cells using a FACSAria III cell sorter (BD Biosciences, La Jolla, CA, USA). For initial selection, the transduced cells were cultured in 2.5 μg/ml puromycin (Gibco) for 5 days, followed by sorting of fluorescent-positive cells using a CytoFlex SRT ( Becman Coulter, Indianapolis, USA). The obtained cell lines (786-O_AREG_GFP, 786-O_FN1_GFP, 786-O_GFP) were cultured in RPMI 1640 Medium (ATCC modification) (Gibco, Life Technologies Ltd, Paisley, UK) supplemented with 10% FBS (Sigma-Aldrich, St. Louis, MO, USA ), and Penimerckcillin–Streptomycin (Biowest, Nuaille, France).

Animal experiments were conducted with the approval of the II Local Ethics Committee in Warsaw (approval no. WAW2/059/2022). NSG male mice (NOD.Cg-Prkdc scid Il2rgtm 1 Wjl/SzJ) were purchased from Jackson Laboratories and bred at the local animal facility. The mice were housed in cages under controlled specific pathogen-free conditions (22^°^C ±-2^°^C temperature, 50 ± 5% relative humidity, and a 12 h light/dark cycle). The mice received a complete laboratory rodent diet (Zoolab, Sędziszów, Poland) and had free access to water. When the mice reached 6 weeks of age, their shoulder area was subcutaneously injected with 1 × 10^6 of either 786-O_GFP, 786-O_AREG_GFP, or 786-O_FN1_GFP cells mixed 1:1 with Matrigel (Corning® Matrigel® Matrix GFR PhenolRF Mouse; Sigma-Aldrich, Sant Louis, MO, USA) (8 mice per group). The mice were daily monitored and weighed. After 26 days half of the mice in each group were intravenously given 5x10^5 MSCs dyed with Qtracker™ 655 Cell Labeling Kit (cat. no. Q25021MP, ThermoFisher Scientific, Waltham, MA, USA). After additional 6 days the mice were sacrificed, and the tumours were excised, measured and weighed. The numbers of mice and the experimental setup are shown in the **Supplementary Figure S2**. For intravital imaging, mice were anesthetized with a xylazine/ketamine mixture (5 mg and 100 mg per kg body weight, respectively) administered intraperitoneally. Body temperature was continuously monitored using a rectal probe and maintained at 37°C with a feedback-controlled heating pad throughout the procedure. Mice were injected intravenously with 100 μL of Brilliant Violet 421™ anti-mouse CD31 antibody (BioLegend, CA, USA), diluted 1:10 in sterile 0.9% NaCl. Imaging was initiated 5 minutes after antibody injection to allow for vascular labelling and signal stabilization. Images and Z-stacks were acquired at 20× magnification using the IVM-CMS3 system (IVIM Technology, Seoul, Korea). IVIM Studio v23.12.040 software (IVIM Technology) was used for image acquisition and processing.

Protein isolation for Western blot analysis was performed using RIPA buffer as earlier described (Hanusek et al., 2022). Isolation of proteins for proteomic analysis was performed as earlier described (Popławski et al., 2017) with modifications. Briefly 1.6×10^5^ cells were seeded at 25 cm^2^ . After 48h cells were rinsed 3 times with PBS without calcium and magnesium (ThermoFisher Scientific, Waltham, MA,, USA), trypsinized, centrifuged (250g, 5min), rinsed with PBS, and centrifuged (250g, 5min). Finally, the cells were lysed with 100 μl of protein isolation buffer (0.1M Tris pH 7.5, 2% SDS, 0.1M DTT) and boiled for 5 min. Following centrifugation at 14,500 rpm, 5 min, the supernatant was aliquoted and stored at -70^°^C.

RNA isolation was done using GeneMATRIX Universal RNA/miRNA Purification Kit (EURX, Gdansk, Poland) as earlier described (Popławski et al., 2023).

Reverse transcription and qPCR analysis was performed as earlier described (Hanusek et al., 2022). Sequences of primers are given in **Supplementary Table S1**.

For Western blot analysis, 30 ug of protein was resolved on 12% SDS-PAGE, following a wet 1.5h transfer onto PVDF membrane. The membranes were blocked in 5% non-fat dry milk for 1h, washed in TBST and probed with anti-AKR1C3 antibody (cat. no. 11194-1-AP, Proteintech Group, Rosemont, IL, USA, 1: 1000 dilution in 5% non-fat dry milk) o/n at 4^°^C. The next day the membranes were washed 3 times in TBST, and incubated with secondary anti-rabbit antibody (Goat Anti-Rabbit Immunoglobulins/HRP (cat. no. P0448, Agilent, dilution 1:10,000) for 1h, following signal detection using SuperSignal™ West Pico PLUS Chemiluminescent Substrate (cat. no. 34580, ThermoFisher Scientific, Waltham, MA, USA) in accordance with the manufacturer’s instructions. For β-actin detection, the membranes were stripped and re-probed with anti-β-Actin antibody (NB600-501, Novus, 1:320,000) for 1h at RT, with the following incubation with Goat Anti-Mouse Immunoglobulins/HRP (cat. no. P0447, Agilent, Santa Clara, CA, USA) 1:10,000.

Immunohistochemical analysis was done using archived FFPE tissue specimens of clear cell renal cell carcinoma (ccRCC) with the approval of the local Bioethics Committee (approval no. 21/2025 and 1/2025) using IHCeasy AREG Ready-To-Use IHC Kit (Cat No. KHC0808; Proteintech Group, Rosemont, IL, USA) in accordance with the manufacturer’s protocol.

Immunocytochemical analysis of HIF1A was performed using a confocal microscope (LSM 800, Axio Observer.Z1 LSM 800) and ZEN 3.7 software (Zeiss, Oberkochen, Germany). Cells were seeded on #1.5 coverslips and, following fixation in 4% PFA, permeabilization with 0.25% Triton X-100, and blocking of nonspecific binding with 2% BSA in TBST, they were incubated overnight at 4°C with a primary anti-HIF1A antibody (Novus Biologicals, UK) at a 1:100 dilution. After washing, cells were incubated with a secondary antibody conjugated to AF594 (Cell Signaling Technology, Netherlands) at a 1:500 dilution for 1 hour at room temperature, followed by incubation with ATTO488-conjugated phalloidin (Sigma-Aldrich, St. Louis, MO, USA) for 30 minutes and DAPI for 2 minutes. Coverslips were mounted on glass slides using Fluorescent Mounting Medium (DAKO, Agilent, Santa Clara, CA, USA).

Microarray analysis was done as previously described (Bogusławska et al., 2019). Briefly, the quality of RNA from MSCs treated with CM collected from 786-O_AREG_GFP, or 786-O_GFP cells was analyzed using 2100 Bioanalyzer ( Agilent, Santa Clara, CA, USA). Preparation of targets was done using the GeneChip™ WT PLUS Reagent Kit (ThermoFisher Scientific, Waltham, MA, USA), with the following hybridization to the Affymetrix™ HuGene 2.1 ST Array Strips (Affymetrix, Santa Clara, CA, USA). Array scanning, data normalization and analysis involving the Transcriptome Analysis Console (TAC) Software 4.0 (ThermoFisher, Waltham, MA, USA) and Ingenuity Pathway Analysis Software (IPA, QIAGEN Bioinformatics, Hilden, Germany). was done as described elsewhere (Bogusławska et al., 2019). The data were deposited in NCBI/GEO database (accession number GSE306138).

Label-free proteomic analysis was done using nanoHPLC-MS/MS as previously described (Poplawski et al., 2023). The data were processed using MaxQuant v2.1.4.0 and Perseus v2.0.7.0. Data matrix was filtered for proteins with at least 3 valid values in at least one group and then missing values were imputed. Statistical analysis was done using Student’s t-test + permutation-based FDR (false discovery rate), with threshold: q-value < 0.05. The data were deposited to the ProteomeXchange Consortium (Vizcaíno et al., 2014) via the MassIVE repository with the dataset identifier PXD065400.

Statistical analysis was done on data from least three independent biological experiments using t-test (for two groups comparisons) or ANOVA with Dunnett’s multiple comparisons test (for three groups comparisons). P < 0.05 was considered statistically significant.

## Results

### AREG induces growth of RCC tumours in vivo

In our recent study we found that cells derived from advanced ccRCC tumours secrete high amounts of AREG and FN1 that stimulate MSCs migration in vitro (Popławski et al., 2023). To analyse the influence of AREG and FN1 on growth of renal tumours, we selected 786-O cells that secrete low amounts of AREG and FN1 and weakly stimulate MSCs migration (Popławski et al., 2023), and stably transduced them with GFP-tagged vectors expressing AREG and FN1. Expression of the ectopically expressed AREG, FN1 was statistically significantly higher than in cells transduced with control (GFP) vector (**Figure 1A**). Furthermore, ELISA demonstrated that both proteins were secreted in amounts similar to those observed in advanced RCC cells that stimulated MSCs migration (Popławski et al., 2023).

**Figure 1.**
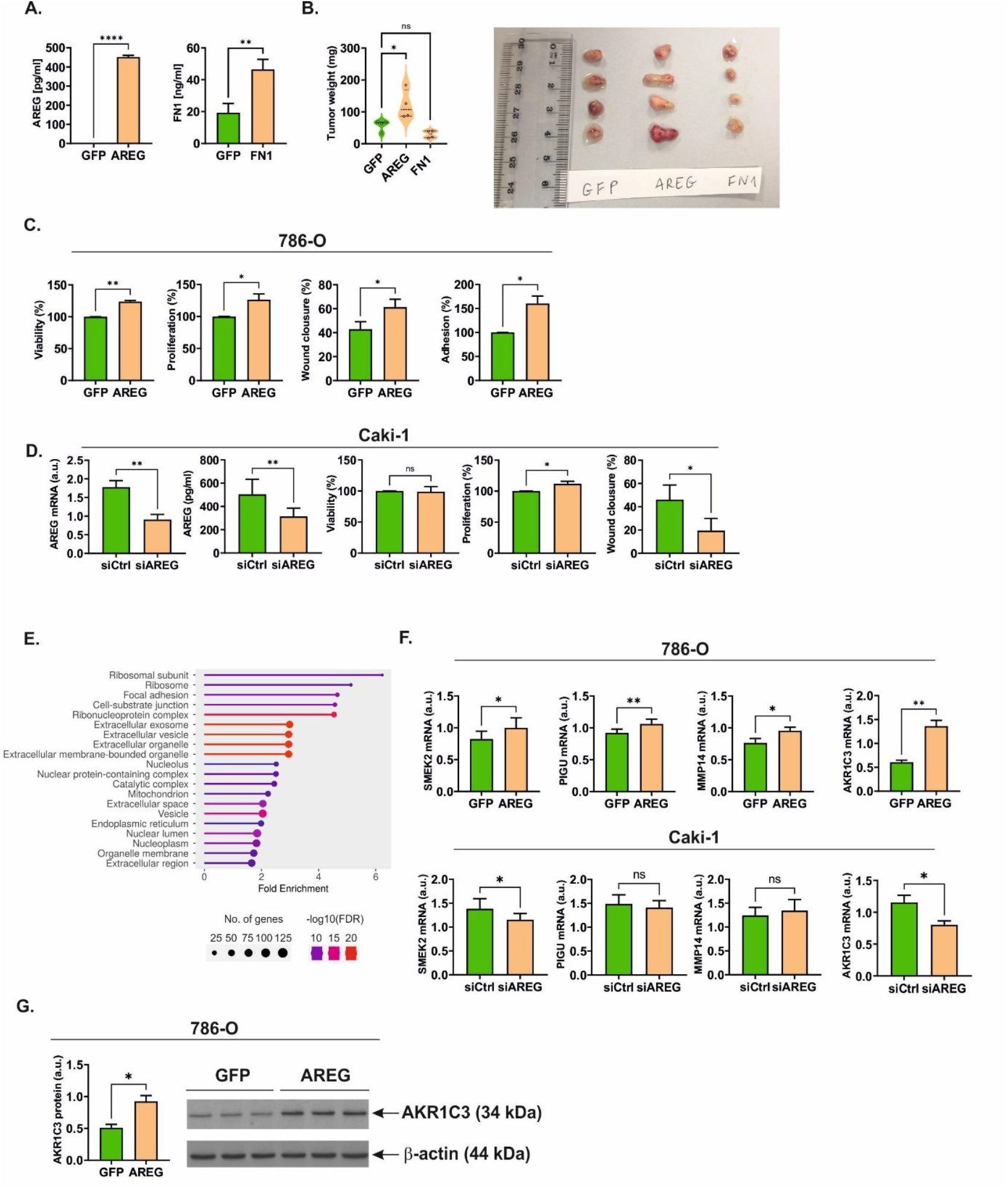
AREG promotes growth of renal tumours in vivo. **A**. Secretion of AREG and FN1 by stably transduced cell lines overexpressing AREG and FN1, respectively. GFP: control vector. **B**. Weight of tumours excised from mice inoculated with 786-O cells overexpressing AREG or FN1. **C**. AREG stimulates viability, proliferation, migration, and adhesion of 786-O cells. **D**. The effects of AREG silencing in Caki-1 cells. **E**. Top enriched cellular components as revealed by the analysis of proteomic data. The analysis was performed using ShinyGO 0.82 (http://bioinformatics.sdstate.edu/go/). **F**. qPCR validation of proteomic data in 786-O_AREG_GFP cells and 786-O_GFP cells, as well as Caki-1 cells with silenced AREG expression (siAREG) when compared with control cells transfected with non-targeting scrambled siRNA (siControl). **G**. Western blot analysis of AKR1C2 in 786-O_AREG_GFP cells and 786-O_GFP cells. The plot shows results of densitometric analysis of 3 independent biological experiments and representative WB scans are shown with each path representing protein isolated from an independent cell culture flask. For raw uncropped WB scans see **Supplementary Figure S4**. The plots show data from at least three independent biological experiments. Statistical analysis was performed using ANOVA with Dunnett’s multiple comparisons test (panel B) or paired t-test (residual panels). * p < 0.05, ** p < 0.01, *** p < 0.001, **** p < 0.0001. In panel (D) wound closure: unpaired t-test: P < 0.05, paired t-test: non-significant.

RCC cells with ectopic expression of AREG, FN1 as well as the respective control cells, were subcutaneously inoculated into the flanks of NSG mice in accordance with the protocol shown in **Supplementary Figure S2**. 32 days following inoculation, mice were killed and tumour weight was evaluated. Ectopic AREG expression increased tumour weight, while FN1 expression did not change growth of xenograft tumours (**Figure 1B**). Furthermore, overexpression of AREG stimulated angiogenesis as revealed by CD31 staining (**Supplementary Figure S3**), and IVIM observations (**Supplementary Movie 1)**.

To explore the mechanisms by which AREG stimulates growth of RCC tumours, we performed functional tests in vitro. AREG overexpression stimulated proliferation, viability, migration, and adhesion of 786-O cells in vitro (**Figure 1C**). Next, we silenced AREG in Caki-1 cells in which AREG is highly expressed (Popławski et al., 2023) (**Figure 1D**). This resulted in approximately 50% reduction of AREG mRNA and moderate suppression of concentration of AREG secreted into the cell culture medium. Viability was not affected, while proliferation was moderately increased by AREG silencing in Caki-1 cells. In contrast, suppression of AREG expression in Caki-1 cells resulted in substantial reduction of migration (**Figure 1D**) which was in sharp contrast to the elevated motility of 786-O cells with AREG overexpression (**Figure 1C**).

These results indicated that increased expression of AREG promotes proliferation, viability, migration and adhesion of ccRCC cells to stimulate growth of ccRCC tumours and enhance angiogenesis.

### AREG reprograms ccRCC proteome

To explore more in depth the mechanisms behind the pro-tumourous AREG effects, we performed proteomic analysis of ccRCC overexpressing AREG (**Supplementary Table S2**). The expression of 343 proteins was altered in 786-0_AREG cells when compared with control cells transduced with empty vector. Notably, GO analysis revealed that the most enriched cellular components included ribosomes, focal adhesion, nucleolus, extracellular space or nucleus (**Figure 1E**). Furthermore, we also found that AREG overexpression in 786-O cells resulted increased expression of the well-known tumor promoters, including SMEK2, PIGU, MMP14, and AKR1C3 (**Supplementary Table S2)**, which was further confirmed by qPCR (**Figure 1G**). In Caki-1 cells with silenced AREG expression we observed moderate suppression of SMEK2 and AKR1C3 mRNA, while expressions of PIGU and MMP14 were not changed (**Figure 1F)**. Strikingly, we observed strong positive correlation between protein levels of AREG and AKR1C3 (r = 0.95, p = 0.046) in the results of proteomic analysis of 786-O_ AREG_GFP cells. In line with this observation, Western blot analysis confirmed that AKR1C3 protein was upregulated in 786-O_ AREG_GFP cells when compared with control cells (**Figure 1G**).

### AREG secreted by RCC recruits MSCs in vivo and reprograms their transcriptome towards stimulated migration and altered immunomodulation

To evaluate the effects of AREG and FN1 on MSCs migration in vivo, mice with ccRCC xenografts were intravenously given labelled MSCs (**Figure 2A**). ICC analysis revealed massive invasion of MSCs in tumours overexpressing AREG while only slight traces of MSCs were found in controls and tumours overexpressing FN1. Intravital microscopy confirmed that within 24h from injection, MSCs invaded AREG-expressing tumours (**Figure 2B, Supplementary Movie 2)**. These results indicated that overexpression of AREG induces migration of MSCs and angiogenesis in vivo.

**Figure 2.**
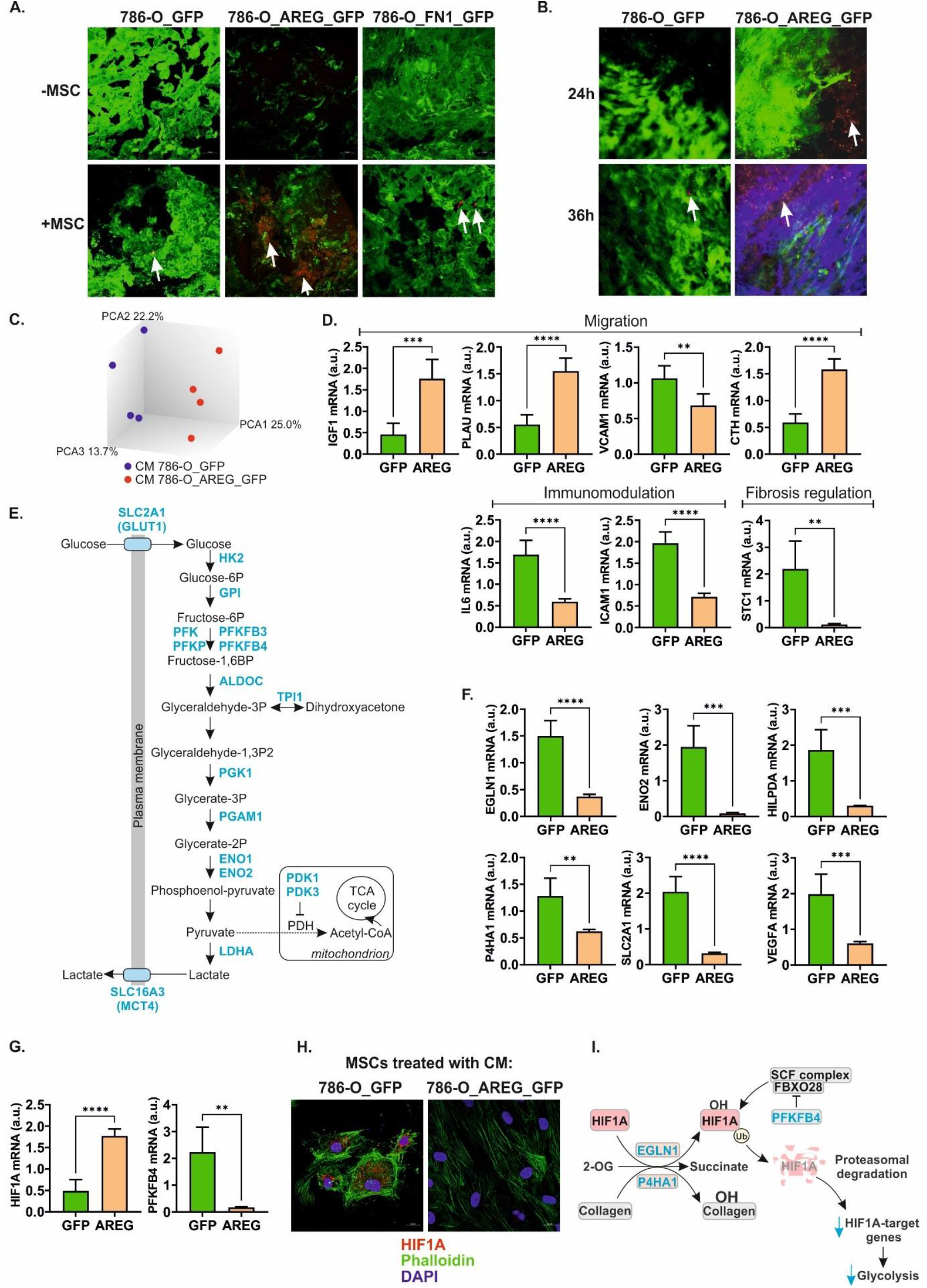
AREG secreted by ccRCC cells reprograms MSCs. **A**. Post-mortem analysis of tumour tissues excised from mice that were given labelled MSC (+MSC) or control mice (-MSC). White arrows point to the MSCs. **B**. Intravital analysis of tumours overexpressing AREG or control tumours, from mice that were given labelled MSCs (red). Green: tumour cells, blue: CD31 (blood vessel marker). **C**. PCA of microarray-analysis of MSCs incubated with CM from 786-O_AREG_GFP or control cells (786-O_GFP). **D**. Altered expression of MSCs genes involved in migration, immunomodulation and regulation of fibrosis. The plots show results of qPCR validation of microarray analysis of MSCs incubated with CM from 786-O-AREG_GFP cells and 786-O-GFP cells (n=5 independent experiments). **E**. A drawing showing that incubation of MSCs with CM from 786-O_AREG_GFP cells globally downregulates genes involved in glycolysis. Blue font: decreased gene expression (microarray data) Please see **Supplementary Table S3** for detailed expression data. **F**. qPCR validation of microarray data – HIF1A targets. Incubation of MSCs with CM from 786-O_AREG_GFP cells triggers HIF1A degradation. The plots show downregulation HIF1A-dependent genes. **G**. CM from 786-O_AREG_GFP cells alters expression of HIF1A and PFKFB4 mRNA in MSCs. **H**. CM from 786-O_AREG_GFP cells triggers HIF1A degradation in MSCs. **I**. The model showing how decreased expression of PFKFB4 acting as a repressor of HIF1A degradation may contribute to the stabilization of HIF1A in MSCs incubated with CM from 786-O_AREG cells. Statistical analysis was performed using paired t-test. * p < 0.05, ** p< 0.01, *** p < 0.001, *** p < 0.0001.

Incubation of MSCs with CM isolated from 786-O_AREG_GFP cells altered expression of 568 genes, including 424 that were upregulated and 144 that were downregulated (**Supplementary Table S3**). PCA showed a clear separation of the two groups, confirming robustness of the analysis (**Figure 2C**). Remarkably, the most upregulated gene was IGF1, known as a stimulator of MSCs migration to injured kidney (Liu et al., 2016; Xinaris et al., 2013). Expression of PLAU and CTH, the two other mediators of MSCs migration (Krstić et al., 2015; Wu et al., 2022) was also increased. In contrast, expression of VCAM1, an inhibitor of MSCs migration (Nishihira et al., 2011) was downregulated (**Supplementary Table S3**, (**Figure 2D**). The most downregulated gene was antifibrotic gene STC1. Furthermore, treatment of MSCs with CM from 786-O_AREG_GFP cells reduced expression of genes involved in immunomodulation, including IL6 and ICAM1, required for immunosuppressive MSCs activities (**Supplementary Table S3, Figure 2D**).

These data indicated that treatment with CM from RCC cells secreting AREG alters expression of MSCs genes towards activation of migration and fibrosis and altered immunomodulation.

### RCC-secreted AREG induces HIF1 degradation and glycolysis inhibition in MSCs

Ingenuity Pathway Analysis showed that glycolysis was among the top canonical pathways altered in MSCs by 786-O_AREG_GFP cells CM (**Table 1, Supplementary Table S3**). The other pathways indicated by IPA included HIF1 signalling, tumour microenvironment pathways, glycolysis I, and apelin endothelial signalling pathway (**Table 1, Supplementary Table S3**). Indeed, microarray data revealed global downregulation of glycolytic genes (**Figure 2E**) and HIF1A targets (**Figure 2F)**. In agreement with these findings, HIF1A emerged is the top upstream regulator predicted to be inhibited (**Table 2**).

**Table 1.** Top canonical signalling pathways altered in MSCs incubated with CM from AREG-overexpressing 786-O RCC cells. The table shows results of IPA analysis.

| Name | p-value | Overlap |
| --- | --- | --- |
| HIF1 Signaling | 1.23E-13 | 15.2 % |
| Tumour Microenvironment Pathway | 8.01E-13 | 15.7 % |
| Glycolysis I | 8.00E-10 | 41.7 % |
| Apelin Endothelial Signaling Pathway | 3.27E-08 | 13.9 % |
| Thyroid Cancer Signaling | 5.67E-08 | 18.4 % |

**Table 2.** Top predicted upstream regulators in MSCs incubated with CM from AREG-overexpressing 786-O cells. The table shows results of IPA analysis.

| Name | p-value | Predicted Activation |
| --- | --- | --- |
| dimethyl n-oxalyl-glycine | 9.36E-62 | Inhibited |
| HIF1A | 9.10E-53 | Inhibited |
| cobalt chloride | 2.02E-39 | Inhibited |
| EGLN | 8.45E-31 | Activated |
| dactolisib | 8.46E-27 | Activated |

Since HIF1A mRNA expression was increased (**Supplementary Table S3, Figure 2G**), this suggested that inhibition of its activity resulted from posttranslational regulation. Incubation of MSCs with CM from 786-O_AREG_GFP cells substantially reduced expression of PFKFB4 (6-phosphofructo-2-kinase/fructose-2,6-biphosphatase 4), a regulator of ubiquitination and proteasomal degradation of HIF1A (**Supplementary Table S3, Figure 2G)**. PFKFB4 depletion leads to enhanced HIF1 degradation (Phillips et al., 2022). In line with these findings, ICC analysis revealed that expression of HIF1A was decreased in MSCs incubated with CM from 786-O_AREG_GFP cells (**Figure 2H)**.

To further validate the impact of AREG on hypoxia-induced signalling pathways, we compared transcriptomic changes induced by 786-O_AREG_GFP cells CM with those induced in MSCs by hypoxia (Zielniok et al., 2021). We found 107 common genes that were altered in both datasets; interestingly, the expression of genes in MSCs incubated with CM from 786-O_AREG_GFP cells was changed in the opposite direction when compared with MSCs incubated with hypoxia (**Supplementary Table S3**). This observation confirmed that treatment of MSCs with CM from ccRCC overexpressing AREG reverses changes induced in MSCs by hypoxia.

Altogether, those analyses confirmed that incubation of MSCs with CM from ccRCC cells overexpressing AREG induces HIF1A degradation and global suppression of genes regulated by hypoxia signalling pathway culminating in glycolysis shut down (**Figure 2I)**.

### AREG is overexpressed by advanced human RCC tumours

To validate associations of AREG with ccRCC pathology, we evaluated its expression in human samples. IHC confirmed that AREG was expressed in RCC tumours, with more intense signals coming from advanced tumours (**Figure 3A, Supplementary Figure S4**).

**Figure 3.**
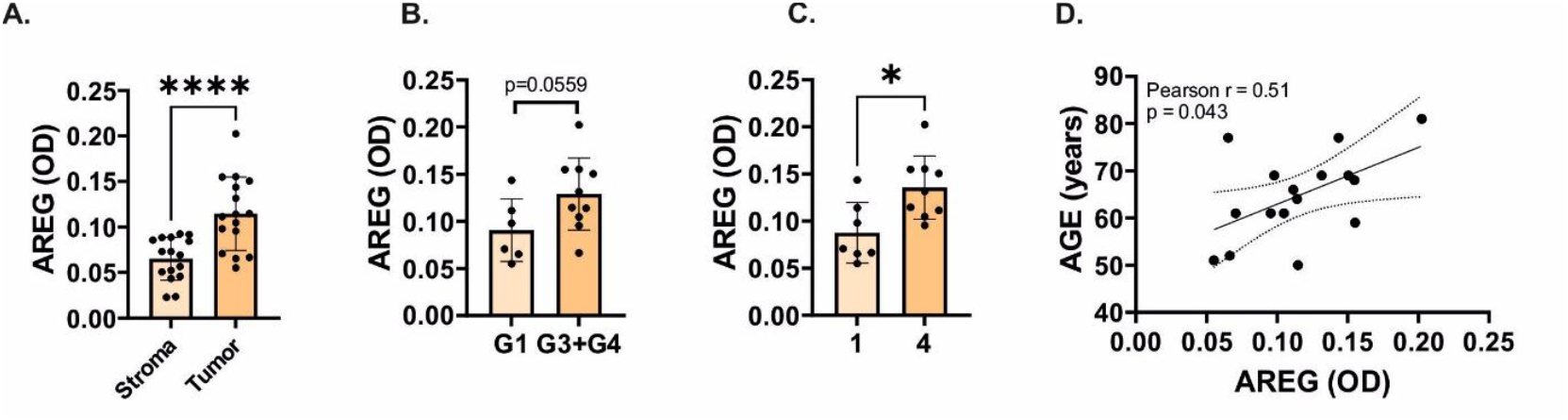
Increased AREG expression in human RCC tumours. The results of IHC analysis of AREG protein in RCC tumours. The plots show mean optical density (OD = log10(max_intensity/mean_intensity)) obtained from the 5 randomly picked microscopic fields per each tissue sample. The analysis was performed on tissue samples from 16 patients. **A**. AREG expression in matched-paired areas of tumour (n=16) and stroma (16). **B**. AREG expression in ccRCC tumours of lower grade (G1, n=6) when compared with tumours of higher grade (G3+G4, n=10). **C**. AREG expression in ccRCC tumours classified as TNM Stage 1 (n=7) and Stage 4 (n=9). Statistical analysis was performed using t-test. * p < 0.05. **** p < 0.0001. **D**. AREG expression in tumours correlates with age of patients. See also **Supplementary Table S4** (patients clinical data) and **Supplementary Figure S4** (microscopic images of IHC).

## Discussion

The physiological role of MSCs is to support wound healing. This process is highly stimulated when MSCs are conditioned under hypoxia (Mathew et al., 2017; Riis et al., 2017; Y. Yang et al., 2022). Tumours, often referred to as ‘wounds the do not heal’, are characterized by highly hypoxic milieu, which raises a puzzle of how MSCs’ healing properties are prevented under oxygen-depriving conditions. Our study provides answer to this jigsaw. Here, we show a novel mechanism by which AREG secreted by renal cancer cells acts in autocrine and paracrine manner to stimulate tumour growth. In MSCs, ccRCC-derived AREG triggers HIF1A degradation and suppression of hypoxia-regulated genes, including those involved in glycolysis, immunomodulation and fibrosis inhibition, as well as activation of genes involved in migration, supporting MSCs homing to ccRCC tumours. In ccRCC cells, AREG promotes viability, proliferation, and migration by upregulating tumour-promoting genes including AKR1C3, SMEK2, PIGU, and MMP14. Altogether, those changes culminate in enhanced angiogenesis and growth of ccRCC tumours in vivo.

Amphiregulin (AREG) is a multifunctional protein, primarily recognized as the EGFR ligand. However, relative to the major EGFR ligand, the EGF, AREG is considered as a weak activator. AREG is widely associated with metastatic properties of cancer cells, as well as chemo- and radioresistance (Bordoli et al., 2011). It also promotes cancer cells proliferation, migration, invasion and evasion of apoptosis. AREG expression in tumour tissues and/or serum positively correlates with poor prognosis for lung, breast, liver, colon, and prostate cancers (Bordoli et al., 2011).

The primary goal of our study was to identify mechanisms by which AREG released by ccRCC cells stimulates migration of MSCs. We found that incubation of MSCs with CM derived from ccRCC cells overexpressing AREG induced expression of several powerful regulators of MSC migration, including. IGF1 (Insulin-Like Growth Factor 1; the most upregulated gene), CTH, and PLAU. It also decreased expression of VCAM1, an inhibitor of MSCs migration. IGF1 is an important regulator of MSCs proliferation, self-renewal, pluripotency and differentiation (Youssef et al., 2017). Moreover, IGF1 is the key stimulator of MSCs’ migration to injured kidney (Liu et al., 2016; Xinaris et al., 2013), making it a perfect mediator of an AREG-induced MSCs homing to ccRCC tumours. Interestingly, MSCs-derived IGF1 stimulates proliferation and protects against apoptosis protection proximal tubule cells (Imberti et al., 2007; Liu et al., 2016). It also protects proximal tubules against cisplatin-induced cytotoxicity (Imberti et al., 2007). All this data suggests that IGF1 secreted by MSCs in response to AREG released by cancer cells reciprocally supports growth of ccRCC tumours. Furthermore, in lung cancer AREG activates IGF1 receptor to induce secretion of IGF1 and attenuate apoptosis (Gigić et al., 1975). This suggests that IGF1 overexpression in MSCs exposed to AREG derived from ccRCC cells could be mediated by IGF1R. Another MSCs gene upregulated in response to CM from 786-O_AREG_GFP cells was CTH which encodes cystathionine gamma-lyase, involved in conversion of L-cystathionine into L-cysteine, and the following production of hydrogen sulphide (H2S), a crucial gaseous signalling molecule. H2S is involved in stimulation of MSCs migration. HMGB1-induced CTH expression leads to increased H2S synthesis, activation of calcium channel, extracellular calcium efflux and enhanced MSCs migration (Wu et al., 2022). PLAU (also known as UPA) encodes plasminogen activator, a secreted protease that converts plasminogen to plasmin which in turn induces ECM remodelling catalysed by matrix metalloproteinases. PLAU is also capable of activating intracellular signalling pathways. PLAU overexpression stimulates MSCs migration to tumours cells, including those derived from prostate and breast cancers and glioma (Pulukuri et al., 2010), while extracellular AREG stimulates PLAU production by breast epithelial cells (Giusti et al., 2003) which fits our observations in ccRCC cells. VCAM1 is an inhibitor of MSCs migration and its knock-down increases motility of BM-MSCs (Nishihira et al., 2011), while in our study VCAM1 expression was decreased in response to CM from 786-O_AREG_GFP cells. Altogether, these data indicate that AREG secreted by ccRCC cells alters expression of migration regulators (IGF1, CTH, PLAU, and VCAM1), concomitantly increasing the recruitment of MSCs to ccRCC tumours.

MSCs have strong immunomodulatory properties (Weiss and Dahlke, 2019). Several previous studies demonstrated that immunoregulatory properties of MSCs are tightly associated with glycolytic switch (Contreras-Lopez et al., 2021, 2020; Jitschin et al., 2019; Liu et al., 2019). In particular, HIF1 inhibition switches MSCs metabolism from glycolysis to OXPHOS and reduces their ability to produce immunoregulatory molecules including ICAM, IL-6, and NO. This finally attenuates the ability of MSCs to inhibit generation of TH1 and Th17 cells (Contreras-Lopez et al., 2020). Furthermore, the immunoregulatory properties of MSCs at glycolytic switch depend on AMPK activity (Contreras-Lopez et al., 2021). In our study, incubation of MSCs with 786-O_AREG_GFP CM decreased expression of PRKAA2, encoding the catalytic subunit of AMPK. In line with these findings, we observed global downregulation of genes encoding glycolytic enzymes, GLUT1 transporter, as well as PDK1 and PDK3, inhibitors of pyruvate dehydrogenase complex in MSCs incubated with CM from ccRCC cells overexpressing AREG. All these changes were associated with decreased expression of ICAM1 and IL6. It is known that extracellular AREG reduces production of iNOS, IL-6 and IL-1b in microglia (Prasad et al., 2023), while IL6 is a target gene for HIF1A (Fan et al., 2023) which further strengthens the validity of our findings. Silencing of IL6 in MSCs reduces their immunosuppressive properties, in particular their ability to suppress proliferation of activated T-cells (Dorronsoro et al., 2020). Prostaglandin E2 is one of key factors mediating the ability of BM-MSCs to inhibit proliferation and cytokine production by T CD8+ cells (Li et al., 2014). In our study, MSCs incubated with CM from 786-O_AREG_GFP cells reduced expression of PTGES (**Supplementary Table S3**), encoding the terminal enzyme of the PGE2 biosynthesis pathway. Interestingly, hypoxia induces PTGES expression in MSCs (Kojima et al., 2019) which further confirms that the changes induced in AREG-treated cells may result from HIF1A-degradation. The other changes in MSCs transcriptome that reflect HIF1A degradation are reduced expressions of NDUFA4L2 and MIR210HG (**Supplementary Table S3**). Under hypoxia, expressions of NDUFA4L2 and miR-210 are activated to attenuate activity of mitochondrial ETC complexes (Lee et al., 2020). All this data suggest that in response to AREG released by ccRCC cells, MSCs attenuated their immunoregulatory properties. Remarkably, these changes in immunoregulatory profiles of MSCs are in line with the unique immune features of ccRCC tumours, in particular high infiltration by T CD8+ lymphocytes (Becht et al., 2016; Borcherding et al., 2021; Giraldo et al., 2019).

The association between MSCs glycolytic suppression and reduced immunoregulation were so far reported in the context of non-neoplastic diseases such as Delayed-Type Hypersensitivity, Graft versus Host Disease or inflammation (Contreras-Lopez et al., 2021, 2020; Jitschin et al., 2019; Liu et al., 2019). To our knowledge, this is the first study showing that glycolytic switch in MSCs can also be induced by cancer.

There are several mechanisms that may contribute to HIF1A degradation in MSCs. Incubation of MSCs with CM from AREG overexpressing ccRCC cells resulted in reduced expression of P4HA1, a prolyl 4-hydroxylase and a key enzyme in collagen synthesis. P4HA1 contributes to HIF1A protein stabilization and its silencing facilitates HIF-1α protein ubiquitination and degradation in an oxygen-independent manner (Xiong et al., 2018). Ubiquitination and degradation of HIF1A is triggered by its prolyl hydroxylation catalysed by EGLN1 (alias PHD2, Hypoxia-Inducible Factor Prolyl Hydroxylase 2). In this reaction EGLN1 utilizes alpha-ketoglutarate (aKG) as a co-substrate that is rapidly converted to succinate. During collagen hydroxylation, P4HA1 acts in a similar way, utilizing aKG and converting it to succinate. This way, P4HA1 reduces availability of aKG for EGLN1, resulting in decreased HIF1A hydroxylation and degradation. Silencing of P4HA1 leads to rapid increase of aKG, facilitating EGLN1-mediated HIF1A hydroxylation and degradation (Xiong et al., 2018). However, in our study, expression of both P4HA1 and EGLN1 was reduced in MSCs incubated with CM from c786-O_AREG_GFP cells, suggesting that HIF1a degradation could be mediated by other mechanism(s). Indeed, incubation of MSC with CM containing AREG resulted in substantial downregulation of PFKFB4 (6-Phosphofructo-2-Kinase/Fructose-2,6-Biphosphatase 4), a multifunctional protein known for its role in the interconversion of glycolytic by products fructose 6-phosphate (F6P) to fructose 1, 6-diphosphate (FBP). Remarkably, PFKFB4 depletion in glioma cells leads to enhanced HIF1 degradation and suppression of glycolytic genes (Phillips et al., 2022). Thus, the loss of PFKFB4 in MSCs incubated with CM derived from ccRCC cells overexpressing AREG, may lead to enhanced loss of HIF1A, and result in decreased expression of HIF1A-dependent genes. PFKFB4 acts by repressing the activity of FBXO28 (F-box only protein 28), a ubiquitin E3 ligase that ubiquitinates HIF1A and targeting it for proteasomal degradation (Phillips et al., 2022). Interestingly, in comparison to other types of cells and tissues, MSCs show the highest level of FBXO28 expression (**Supplementary Figure S5**), suggesting that this pathway of HIF1A degradation in MSCs is plausible. Furthermore, expression of FGF11 that stabilizes HIF1A upstream of proteasomal degradation (Lee et al., 2017) was reduced in MSCs following CM incubation. In contrast to decreased protein, the expression of HIF1A mRNA was increased in MSCs incubated with CM from 786-O_AREG_GFP cells. This may possibly result from decreased PRKAA2 expression. PRKAA2 encodes a catalytic subunit of the AMP-activated protein kinase (AMPK). Silencing of PRKAA2 leads to upregulation of HIF1A mRNA in hepatoblastoma cells (Xie et al., 2023).

Apart from genes involved in MSCs migration, glycolysis and immunoregulation, incubation of MSCs with CM from 786-O_AREG_GFP cells affected also other genes crucial for MSCs functioning. The most downregulated gene was STC1 encoding stanniocalcin-1, a multifunctional glycoprotein that is secreted by MSCs. Similar to other genes analysed in this study, STC1 expression is regulated by HIF1A (Law et al., 2010; Yeung et al., 2005). STC1 has strong antifibrotic properties. Treatment of proximal tubules with recombinant STC1 reduces TGFB1 induced fibrosis (E. M. Yang et al., 2022). Moreover, MSCs that migrate to the injured lungs attenuate fibrosis by secreting STC1 (Ono et al., 2015). Fibrosis is associated and contributes to the progression and therapy resistance in RCC (J.-Y. Chen et al., 2023). We found that MSCs incubated with CM from AREG-overexpressing ccRCC cells dramatically reduced the expression of STC1, fibrosis inhibitor. This suggests that ccRCC cells reduce antifibrotic capacities of MSCs by AREG-mediated STC1 downregulation.

We hypothesize that ccRCC tumours may ‘educate’ MSCs to adapt to the harsh tumour microenvironment. Solid tumours are highly hypoxic and serum deprived. Both hypoxia and serum deprivation induce MSCs apoptosis (Zhu et al., 2006). This may suggest that in advanced ccRCC tumours, increased secretion of AREG allows reprogramming of MSCs to shut-down HIF1A signaling and reduce immunoredulatory MSCs activities. This hypothesis is further supported by the fact that incubation of MSCs with AREG-rich CM induced expression of CTH, encoding a cystathionine gamma lyase, an enzyme that protects MSCs against hypoxia and serum deprivation (Guo et al., 2015).

Our study shows that apart from its effect on MSCs, AREG has profound impact on ccRCC tumour growth and angiogenesis. ccRCC tumours express EGFR (Minner et al., 2012), indicating that amphiregulin may act in an autocrine manner to promote ccRCC progression. The stimulatory effect of AREG on ccRCC viability, proliferation, migration, and tumour growth is consistent with its established role as an oncogenic protein (Busser et al., 2011). We also observed that tumours overexpressing AREG were enriched in vessels. This is consistent with previous findings showing that endothelial tube formation can be blocked by anti-AREG antibodies (Bordoli et al., 2011).

AREG overexpression in ccRCC cells altered expression of multiple cancer-related proteins, including SMEK2, a regulatory subunit of serine/threonine protein phosphatase 4 (PP4) (Raja et al., 2022; Zhu et al., 2024) or MMP14 (aka MT-MMP), a membrane-bound metalloproteinase that digests components of extracellular matrix, activates proMMP2 and promotes cancerous invasion (Knapinska and Fields, 2019). MM14 is the major mediator of RCC invasiveness (Petrella and Brinckerhoff, 2006) and MMP14 transfection into non-malignant kidney epithelial cells leads to their cancerous transformation and invasive tumours formation in mice (Soulié et al., 2005). In line with these findings, high MMP14 expression correlates with poor prognosis for RCC patients (Zhao et al., 2022). Finally, we found that overexpression of AREG in ccRCC cells enhanced expression of AKR1C3, a 17-ketosteroid reductase that converts delta4-androstenedione to testosterone. AKRC1C3 is recognized as an oncogenic protein that is broadly upregulated in various cancer types and promotes cancerous proliferation, migration, angiogenesis, and radio- or chemoresistance (Dozmorov et al., 2010; Huang et al., 2025; Li et al., 2024). It is also a target of several anticancer drugs that are under tests in preclinical and clinical settings (Li et al., 2024). In the context of our study, it is interesting that was demonstrated that treatment of human keratinocytes with AREG induces AKR1C3 expression (Vogeley et al., 2022).

Our study opens several important questions. Firstly, the causes of increased AREG expression in advanced ccRCC tumours are unknown. The possible mechanisms may involve transcriptional activation (e.g. mediated by c-JUN or p53 that directly promote AREG transcription (Hino et al., 2022; Taira et al., 2014)), cytokines (e.g. TGFB1, (Chen et al., 2018)) or microRNAs (e.g. miR-34c-5p (Tung et al., 2017)). Secondly, AREG’s impact on other components of ccRCC TME (e.g. immune cells, CAFs, adipocytes) requires exploration. Finally, the limitation of our study was that we did not analyse RCC-MSCs interaction in co-culture experiments. The reason for this was that we wished to selectively analyse the impact of ccRCC secretome on MSCs migration. However, this kind of experiment does not allow to explore the reciprocal communication between MSCs and ccRCC. Thus, further studies are needed to find how the AREG-reprogrammed MSCs influence the behaviour of ccRCC tumours.

In conclusion, we show that AREG overexpressed and released by ccRCC cells facilitates tumour growth by acting in autocrine and paracrine manner. In MSCs, AREG triggers HIF1A degradation and transcriptional reprogramming, leading to glycolytic switch, increased migration and attenuation of immunoregulatory profile. In ccRCC cells, AREG stimulates expression of oncogenic proteins, including SMEK2, MMP14 and PIGU, leading to increased tumour growth and angiogenesis. Further studies are needed to explore AREG effects on other components of ccRCC TME, including CAFs, immune cells or adipocytes. Finally, our study indicates that AREG could be an attractive target for ccRCC therapy.

## Acknowledgements

The study was financially supported by National Science Center, Poland, grants no. 2018/29/B/NZ5/01211 and 2019/35/B/NZ5/00695. The funding body played no role in the design of the study and collection, analysis and interpretation of data and in writing the manuscript.

## Author contributions

**Conceptualization:** A.P.-W., P.P., **Methodology:** A.P.-W., P.P., T.S. D.N., A.B. (Anna Burdzińska), **Software:** M.K. H.K, **Validation:** P.P., M.G., B.R., A.B (Alex Białas), **Formal analysis:** P.P., M.G., K.B., H.K., M.K., A.B., W.Z.-W., M.P., A.P.W. **Investigation:** P.P., M.G., T.S., G.H., K.B., D,N., H.K., R.I.-N., M.K., U.J., B.S.-R., A.B. (Anna Burdzińska), W.Z.-W., M.P., A.L., M.B., J.Ż-K., N.R., B.R., A.B. (Alex Białas), K.D.A., M.M., A.P.-W. **Resources:** T.S., D.N., M.K., U.J., M.P., J.B., E.S., M.M., W.B., L.P., A.P.-W. **Data Curation:** P.P., M.K., U.J., A.P.-W. **Writing - Original Draft Writing:** A.P.-W. **Review & Editing:** P.P., M.G., G.H., K.B., D.N., H.K., R.I.-N., M.K., U.J., B.S.-R., A.B. (Anna Burdzińska), W.Z.-W., M.P., A.L., M.B., J.Ż.-K., J.B., N.R., E.S., B.R., A.B. (Alex Białas)., K.D.A., M.M., W.B., L.P., A.P.-W. **Visualization: P.P**., **M.G**., **K.B**., **G.H**., **H.K**., **M.K. M.P**., **A.P.W. Supervision:** T.S.,M.K., U.J., J.B., E.S., W.B., L.P., E.S. **Project administration:** A.P.-W. **Funding acquisition:** A.P.-W.

## Material and correspondence

Correspondence and material requests should be addressed to Agnieszka Piekiełko-Witkowska

## Data availability statement

Proteomic and microarray data have been deposited at MassIVE repository (identifier PXD065400) and NCBI/GEO (accession number GSE306138), respectively.

