## Supplementary Data and Figures for "Renal cancer cell-derived amphiregulin recruits mesenchymal stromal cells, induces their glycolytic switch, and promotes tumour growth"

**Supplementary Tables**

**Supplementary Table S1. Primers used in the study.**

**Supplementary Table S2. The results of proteomic analysis of 786-O_AREG_GFP cells and control 786-O_GFP cells.** N=4. The data were deposited to the ProteomeXchange Consortium (Vizcaíno et al., 2014) via the MassIVE repository with the dataset identifier PXD065400.

**Supplementary Table S3. The results of microarray analysis of MSC incubated with conditioned medium retrieved from 786-O_AREG_GFP when compared with 786-O_GFP.** N=4. The data were deposited in NCBI/GEO database (acc. no GSE306138).

**Supplementary Table S4. Clinical data of ccRCC patients whose tumours were analyzed by IHC.**

**Supplementary Figures**

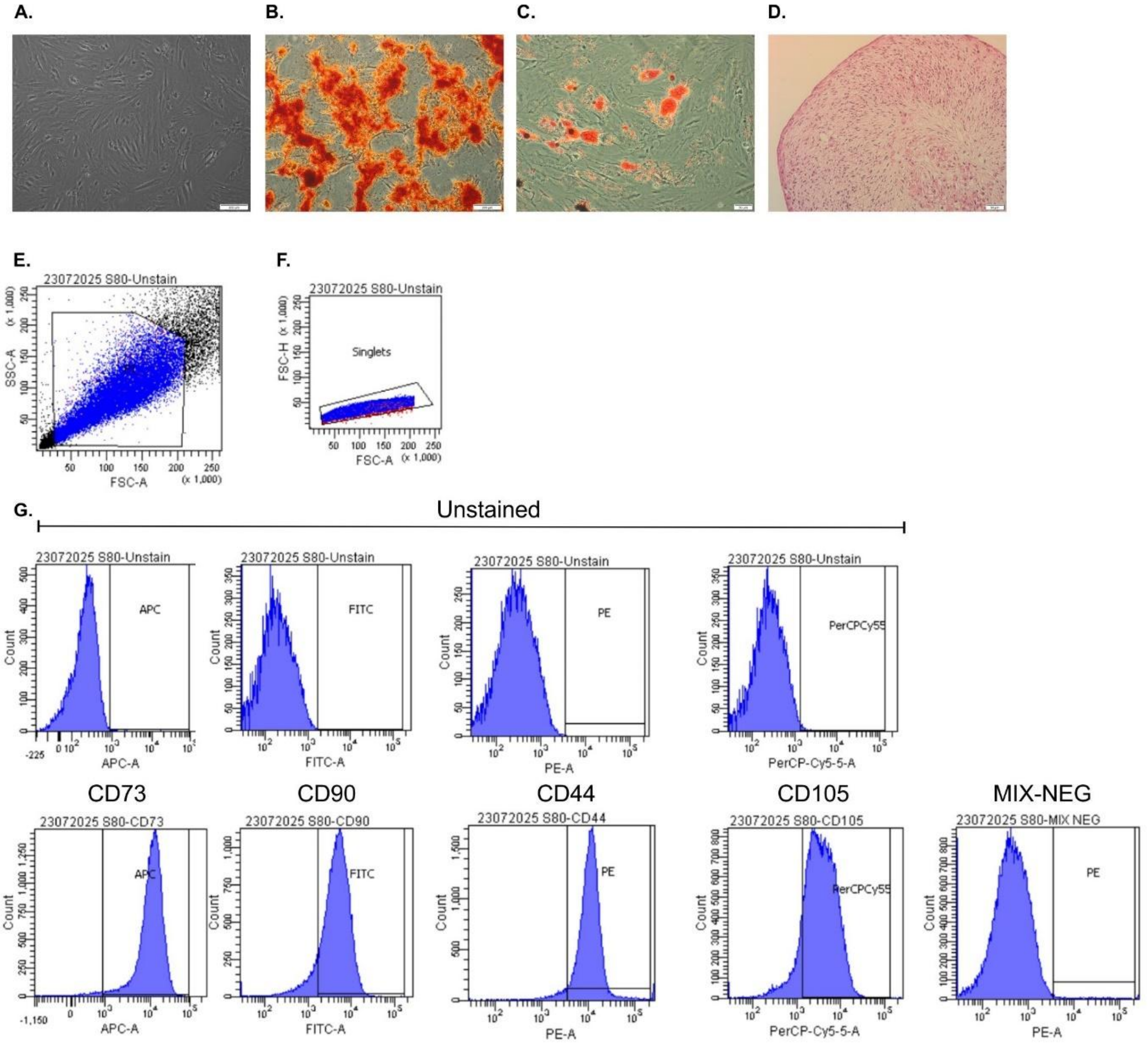


**Supplementary Figure S1. Differentiation and cytometry analysis of MSCs used in the study. A.** Undifferentiated MSCs. **B.** Osteogenic differentiation confirmed by detection of calcium deposits by Alizarin Red staining. **C.** Adipogenic differentiation confirmed by detection of lipid droplets by Oil Red O. **D.** Chondrogenic differentiation confirmed by detection of chondropellet by HE staining. Scale bars: 200 µm (A, B), 20 µm (C), 50 µm (D). **E.** Initial gating: FSC-A vs SSC-A (selection of viable cells based on size and granularity). **F.** Doublet exclusion: FSC-A vs FSC-H (control if only single cells are analyzed). **G.** Marker analysis: Flow-cytometric evaluation of MSCs surface antigens: CD90, CD105, CD73, CD44 and MIX negative (CD34, CD45, CD11b, CD19 and HLA-DR).

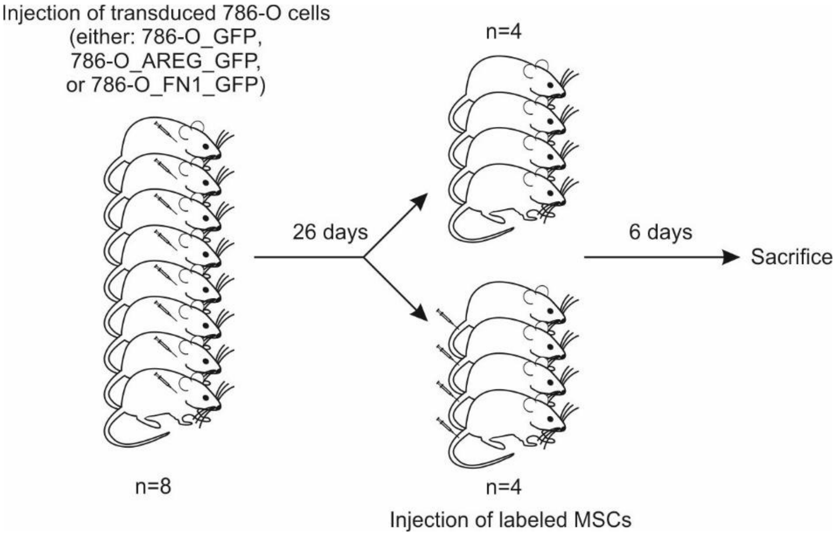


**Supplementary Figure S2. The scheme of mice experiments.** 24 mice were split into three groups (8 mice per group) that were subcutaneously injected with either 786-O_GFP, 786-O_AREG_GFP, or 786-O_FN1_GFP cells (1x10^6 cells per mice). The mice were daily monitored and weighed. After 26 days half of the mice in each group were given 5x10^5 MSCs labelled with Qtracker. After additional 6 days the mice were sacrificed, and the tumours were excised, measured and weighed.

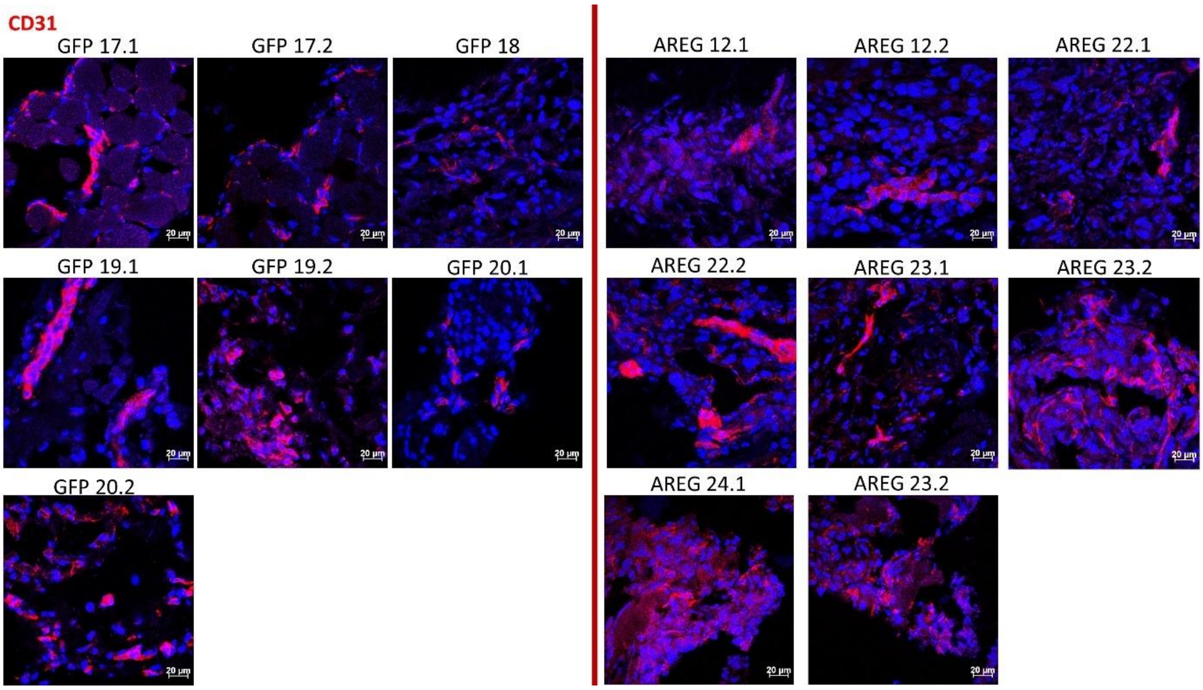


**Supplementary Figure S3. Increased CD31 staining in tumours expressing AREG.** GFP: control tumours (xenografts of 786-O_GFP cells), AREG: tumours with AREG overexpression (xenografts of 786-O_GFP_AREG cells). Red: CD31. Blue: DAPI.

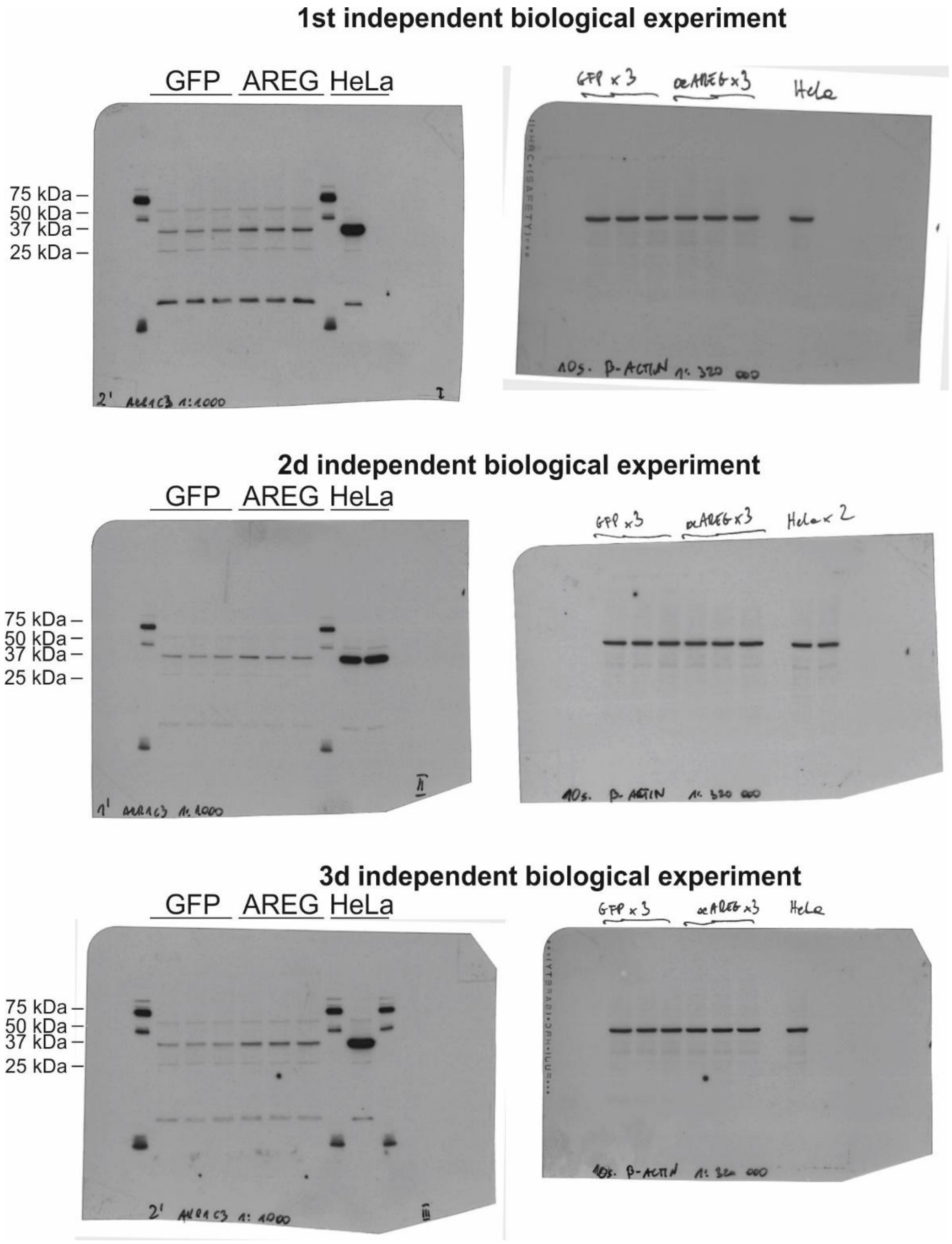


**Supplementary Figure S4. Raw uncropped scan of AKR1C3 WB shown in Figure 1.**

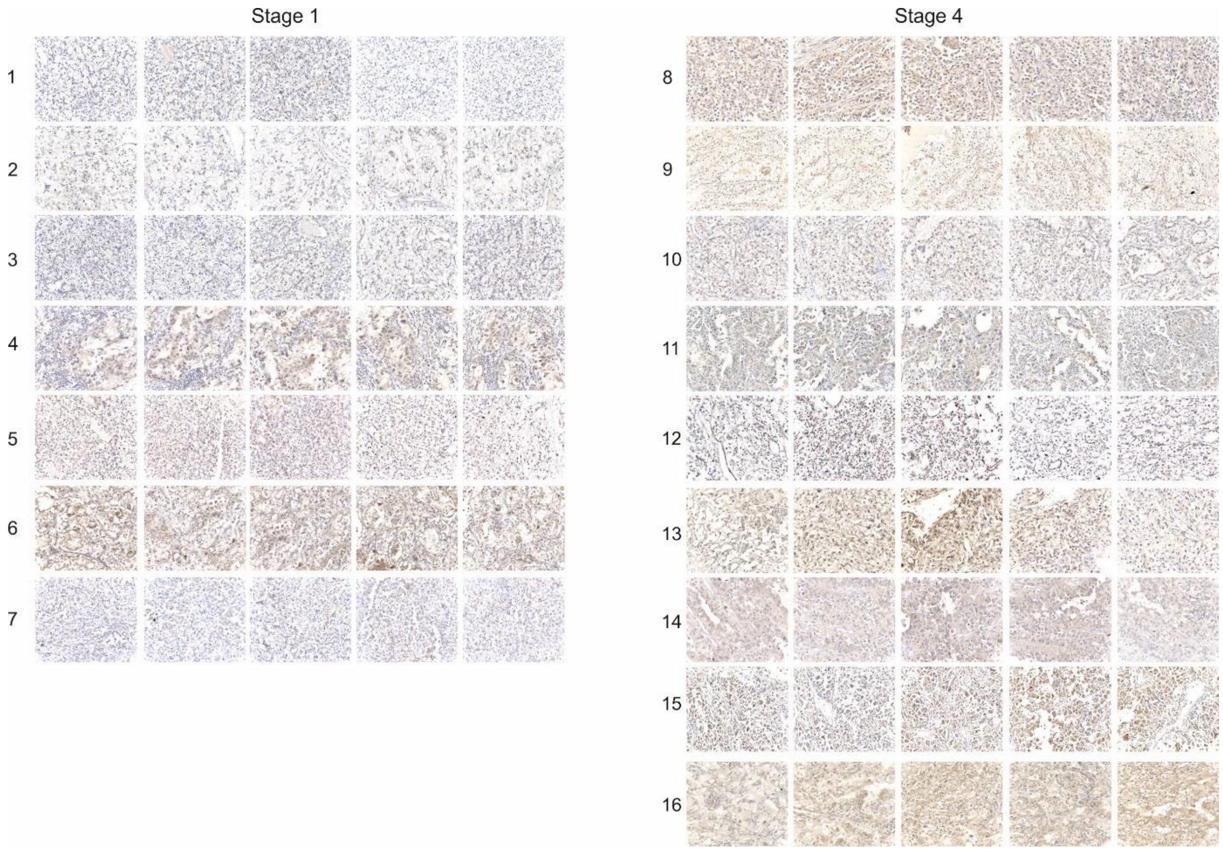


**Supplementary Figure S4. IHC analysis of AREG in ccRCC tumours. For clinical data see Supplementary Table S4.**

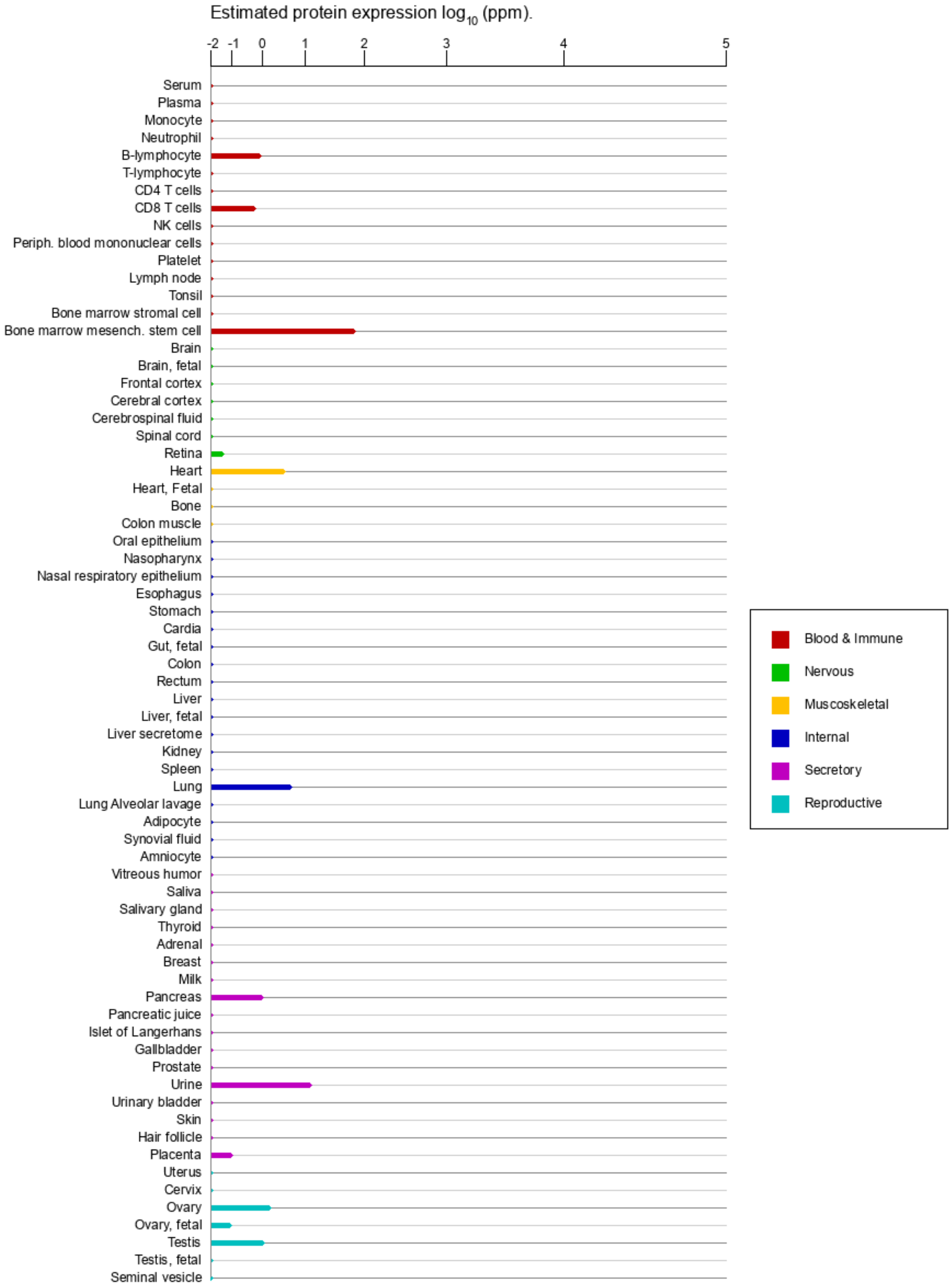


**Supplementary Figure S5. Enhanced expression of FBXO28 in MSC when compared with other tissue and cell types.** The plot was retrieved from GeneCards (https://www.genecards.org/cgi-bin/carddisp.pl?gene=FBXO28) and shows mass spectrometry-based proteomics data integgarted in HIPED (the Human Integrated Protein Expression Database). (Fishilevich et al., 2016).

**Supplementary Movies**

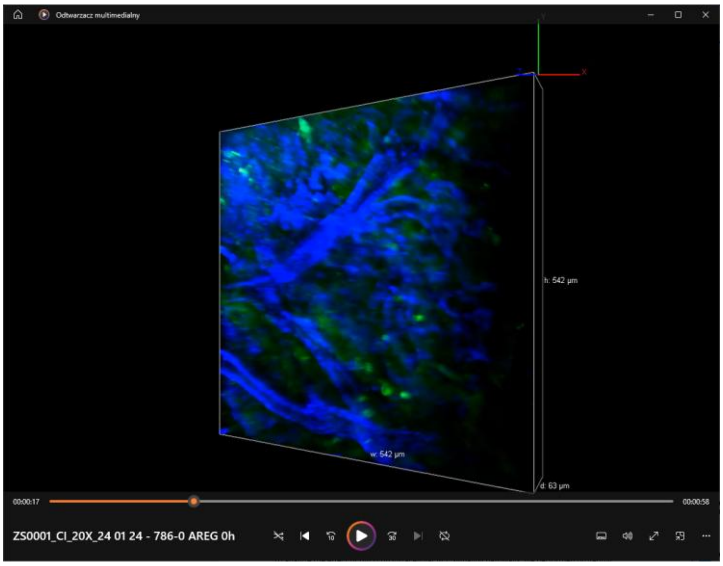


**Supplementary Movie 1.** AREG expression stimulates angiogenesis. The movie shows endothelial cells (blue, anti-CD31 staining) crossing tumour cells (green, GFP).

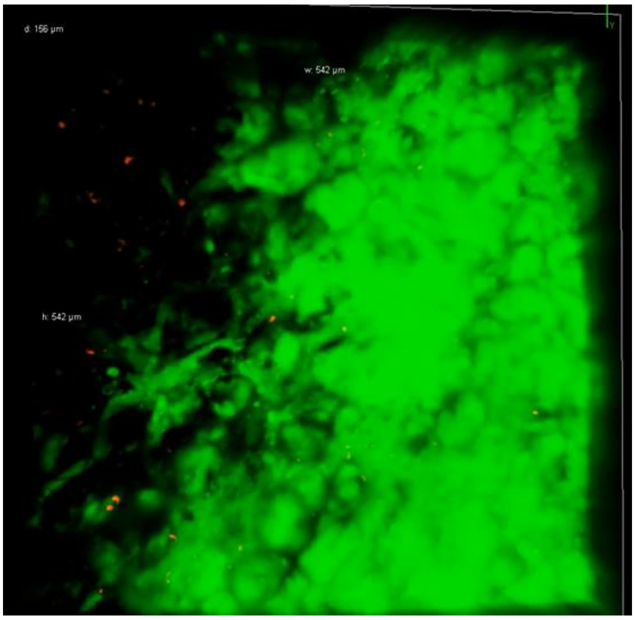


**Supplementary Movie 2. AREG stimulates migration of MSCs toward ccRCC tumours.** The movie shows MSCs (red) approaching ccRCC tumour formed by 786-O-AREG cells (green)
