## Supplementary Table S1 for "Renal cancer cell-derived amphiregulin recruits mesenchymal stromal cells, induces their glycolytic switch, and promotes tumour growth"

**Supplementary Table S1. Primers used in the study.**

| **Target gene** | **Forward** | **Reverse** |
| --- | --- | --- |
| AREG | GTGTCCCAGAGACCGAGTTG | AGGCATTTCACTCACAGGGG |
| CTH | TGGATGATGTGTATGGAGGTACAA | GGGTTTGTGGGGGTTTCGAT |
| EGLN1 | CAAAGCCCAGTTTGCTGACA | TCGTGCTCTCTCATCTGCATC |
| ENO2 | AGCTGGTGAAGGAAGCCATC | ATCGGGAAGGATCAGTGGGA |
| FN1 | GACACATTCCACAAGCGTCA | CTCCAATTTGATAAAACGTCCC |
| HIF1A | CATAAAGTCTGCAACATGGAAGGT | ATTTGATGGGTGAGGAATGGGTT |
| HILPDA | AACCGACTTTCCTCCGGACT | GCTTCATGGCTGAAAGGACC |
| HSPA6 | CGTGCCCGCCTATTTCAATG | AAAAATGAGCACGTTGCGCT |
| ICAM1 | AGTGTGACCGCAGAGGACG | CAGTGGCTGGGCTGGAACC |
| IGF1 | AGAGCCTGCGCAATGGAATA | TTGGGTTGGAAGACTGCTGA |
| IL6 | GGTACATCCTCGACGGCATCT | GTGCCTCTTTGCTGCTTTCAC |
| MMP14 | CCCAACATCTGTGACGGGAA | TTGGTTATTCCTCACCCGCC |
| P4HA1 | TGGAAAATTGACCACAGCACAG | CGTGCAAAGTCAAAATGGGGT |
| PFKFB4 | GGTGAACTTTCAAAGGTGGGC | GAGAGTTGGGCAGTTGGTCA |
| PIGU | CCAGTCTGGCCGAGTTCATT | ACGGAGATACTCCCAAGTCCA |
| PLAU | AACTGCCCAAAGAAATTCGG | TCGGTAAAAGTGACCATTCCC |
| RNA18S1 | CCATCCAATCGGTAGTAGCG | GTAACCCGTTGAACCCCATT |
| SLC2A1 (GLUT1) | GGCTCCTTCTCTGTGG | GGCCAGCAGGTTCATC |
| SMEK2 | TCAGAAACTGAACAGTGTACCA | GGTGCCACAACTGCTTTTCC |
| STC1 | TTCGGAGGTGCTCCACTTTC | CAGGCTTCGGACAAGTCTGT |
| VCAM1 | ACGAACACTCTTACCTGTGC | TCCAAACTCTTGGTTTCCAGG |
| VEGFA | TTGCTCAGAGCGGAGAAAGC | CGTTTAACTCAAGCTGCCTCG |
