## Supplementary Table S4 for "Renal cancer cell-derived amphiregulin recruits mesenchymal stromal cells, induces their glycolytic switch, and promotes tumour growth"

**Supplementary Table S4. Clinical data of ccRCC patients whose tumors were analyzed by IHC.**

| **Patient number** | **Sex** | **Age** | **Grading** | **TNM** | **Cancer stage** |
| --- | --- | --- | --- | --- | --- |
| 1 | K | 51 | G1 | pT1aNx | 1 |
| 2 | K | 52 | G3 | pT1aNx | 1 |
| 3 | M | 61 | G1 | pT1aNx | 1 |
| 4 | M | 64 | G4 | pT1aNx | 1 |
| 5 | M | 69 | G1 | pT1aN0 | 1 |
| 6 | M | 77 | G1 | pT1aNx | 1 |
| 7 | M | 77 | G1 | pT1aNx | 1 |
| 8 | M | 59 | G4 | pT1aNxM1 | 4 |
| 9 | M | 50 | G4 | pT3aN0M1 | 4 |
| 10 | M | 61 | G3 | pT3bN1M1 | 4 |
| 11 | M | 61 | G3 | pT3bNxM1 | 4 |
| 12 | K | 66 | G1 | pT1aNxM1 | 4 |
| 13 | M | 68 | G4 | T3aNxM1 | 4 |
| 14 | M | 69 | G4 | pT3aNxM1 | 4 |
| 15 | M | 69 | G4 | pT2aNxM1 | 4 |
| 16 | K | 81 | G3 | T3aNxM1 | 4 |
